# The constant philopater hypothesis: a new life history invariant for dispersal evolution

**DOI:** 10.1101/023655

**Authors:** António M. M. Rodrigues, Andy Gardner

## Abstract

Life history invariants play a pivotal role in the study of social adaptation: they provide theoretical hypotheses that can be empirically tested, and benchmark frameworks against which new theoretical developments can be understood. Here we derive a novel invariant for dispersal evolution: the “constant philopater hypothesis” (CPH). Specifically, we find that, irrespective of variation in maternal fecundity, all mothers are favoured to produce exactly the same number of philopatric offspring, with high-fecundity mothers investing proportionally more, and low-fecundity mothers investing proportionally less, into dispersing offspring. This result holds for female and male dispersal, under haploid, diploid and haplodiploid modes of inheritance, irrespective of the sex ratio, local resource availability, and whether mother or offspring controls the latter’s dispersal propensity. We explore the implications of this result for evolutionary conflicts of interest – and the exchange and withholding of contextual information – both within and between families, and we show that the CPH is the fundamental invariant that underpins and explains a wider family of invariance relationships that emerge from the study of social evolution.

## Introduction

A number of surprising invariance relationships have emerged from the study of social evolution, whereby a cancelling-out of multiple partial effects of a genetic, ecological or demographic parameter means that it has no net impact upon the evolution of a social behaviour. For example, in the study of sex allocation under “local mate competition” (Hamilton 1967), the number of sons produced by a mother is expected to be independent of her fecundity, in what is known as the “constant male hypothesis” (CMH; Frank, 1985, 1987b; Yamaguchi, 1985). Specifically, the increased extent to which the sons of more-fecund mothers engage in costly competition with male relatives for mating opportunities means that a mother’s proportional investment into sons is expected to be inversely proportional to her fecundity, such that her absolute investment into sons is invariant with respect to her fecundity. Such invariance results provide an important stimulus for scientific advancement. For example, the discovery of the CMH invariant spurred both empirical testing and further development of theory in the field of sex allocation, which has continued in a sustained way from the mid 1980s to the present day (Frank, 1985, 1987a,b,c; May & Seger, 1985; Yamaguchi, 1985; Stubblefield & Seger, 1990; Foster & Benton, 1992; Hasegawa & Yamaguchi, 1995; Petersen & Fischer, 1996; Flanagan et al., 1998; Wool & Sulami, 2001; Ode & Rissing, 2002; Dagg & Vidal, 2004; Akimoto & Murakami, 2012; Akimoto et al., 2012; Rodrigues & Gardner, 2015).

Such invariance results may cross over from their field of origin to illuminate other topics, in which they give rise to new waves of theoretical and empirical research. For example, a surprising discovery that sex ratios are unaffected by the rate of female dispersal – owing to a cancellation of relatedness and kin-competition effects (Bulmer, 1986; Frank, 1986; Taylor, 1988a) – was subsequently shown to translate to the evolution of helping and harming behaviours, stimulating a great deal of further theoretical and empirical study (Taylor, 1992; Wilson et al., 1992; Taylor & Irwin, 2000; Irwin & Taylor, 2001; Perrin & Lehmann, 2001; Gardner & West, 2006; Lehmann et al., 2006; Alizon & Taylor, 2008; El Mouden & Gardner, 2008; Grafen & Archetti, 2008; Johnstone, 2008; Johnstone & Cant, 2008; Kümmerli et al., 2009; Gardner, 2010; Rodrigues & Gardner, 2012, 2013a, 2013b; Yeh & Gardner, 2012). More generally, invariance with respect to transformation is the basis for all analogy and the generalisation of all scientific knowledge to new domains.

Dispersal is a major life history trait and received a considerable amount of attention from both theoreticians and empiricists and has been studied in relation to a variety of factors such as kin competition (Hamilton & May, 1977; Léna et al., 1998; Ronce et al., 1998, 2000; Leturque & Rousset, 2003; Kisdi, 2004; Innocent et al., 2010; Rodrigues & Johnstone, 2014), spatial and/or temporal heterogeneity (Comins et al., 1980; Hastings, 1983; Holt, 1985; Cohen & Levin, 1991; McPeek & Holt, 1992; Gandon & Michalakis, 1999; Leturque & Rousset, 2002; Massol et al., 2010; Rodrigues & Johnstone, 2014), parent-offspring offspring (Motro, 1983; Frank, 1986; Taylor, 1988b; Gandon, 1999; Starrfelt & Kokko, 2010), intragenomic conflict (Farrell et al., 2015), budding dispersal (Gandon & Michalakis, 1999), density-dependent dispersal (Crespi & Taylor, 1990; Travis et al., 1999; Poethke & Hovestadt, 2002; De Meester & Bonte, 2010; Baguette et al., 2011) and other types of condition-dependent dispersal (Ronce, 1998, 2000; Kisdi, 2004; Gyllenberg et al., 2011a,b).

One factor that is likely to have an important impact on the evolution of dispersal is variation in fecundity among group members, i.e. reproductive skew (Vehrencamp, 1983; Hager & Jones, 2009). The social evolutionary consequences of variation in fecundity has received attention in relation to helping and harming behaviour (Frank, 1996; Johnstone, 2008; Bao & Wild, 2012; Rodrigues & Gardner, 2013a) and sex ratio (Yamaguchi, 1985; Frank, 1985, 1987c; Stubblefield & Seger, 1990; Rodrigues & Gardner, 2015). However, the implications for dispersal, and attendant conflicts of interest within and between families, remain to be addressed.

Here we study the evolution of dispersal in groups where the fecundity of breeders vary and report a new invariance result – the “constant philopater hypothesis” (CPH) – in the context of dispersal evolution. We find that, irrespective of variation in maternal fecundity, each mother is expected to make the same absolute investment into philopatric (i.e. non-dispersing) offspring. This is because higher fecundity is associated with one’s offspring facing more stringent kin competition for breeding opportunities when failing to disperse, such that each mother’s proportional investment into philopatric offspring is expected to be inversely proportional to her fecundity. We develop a mathematical kin-selection model to show that the CPH holds for female and male dispersal, under haploid, diploid and haplodiploid modes of inheritance, irrespective of the sex ratio, local resource availability, and whether mother or offspring controls the latter’s dispersal propensity. We provide explicit solutions for variation in resource availability within and between patches, considering both spatial heterogeneity and also temporal heterogeneity for unpredictable and seasonal environments, and we explore the implications of this result for evolutionary conflicts of interest – and the exchange and withholding of contextual information – both within and between families. Finally, we show that the CPH result is the fundamental invariant that underpins and explains a family of other invariance results, including the previously described “constant female hypothesis” (CFH; Frank, 1987c, 1998).

## Model and Results

### Model

We assume an infinite island model (Wright, 1931; Hamilton & May, 1977; Rodrigues & Johnstone, 2014), with *n* mothers in every patch. There are different types of patches, i.e. type-*t* patches with t ∈ T = {1, 2,…, *n*_p_}, and each type differing in its resource availability. Within each patch, each mother is randomly assigned a rank i ∈ I = {1, 2,…, *n*}, and produces a large number of offspring in accordance with her rank, such that no two mothers in the same patch share the same rank, and all mothers sharing the same rank and patch type have the same fecundity. In the asexual version of the model we consider that all offspring are daughters and clones of their mother, and in the sexual version of the model we consider that a fraction *σ*_it_ of the offspring of a rank-*i* mother are sons and a fraction 1-*σ*_it_ are daughters and that there is a haploid, diploid or haplodiploid mode of inheritance. After reproduction, all mothers die, and the offspring of rank-*i* mothers either remain in their natal patch with probability 1-*z*_it_ or else they disperse with probability *z*_it_, with a fraction 1-*c* of dispersers relocating to a new randomly-chosen patch and the remainder *c* perishing en route. We assume that dispersal is controlled either by the offspring themselves or by their mother. In the sexual version of the model, individuals mate at random within their patches following dispersal, with each female mating once, after which all males die. Patches may maintain their resource availability, and therefore remain of the same type, or change their resource availability, and therefore change their type. Females then compete for breeding opportunities, with *n* females being chosen at random within each patch to become the mothers of the next generation, and all other females dying, which returns the population to the beginning of the lifecycle.

### Evolution of dispersal

Applying kin-selection methodology (Hamilton, 1964; Taylor & Frank, 1996; Frank, 1997, 1998; Rousset, 2004; Taylor et al., 2007), we find that an increase in the probability of dispersal of an offspring of a rank-*i* mother in a type-*t* patch is favoured when

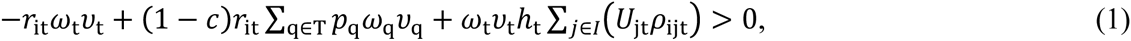

where: *w*_t_ is the probability that an individual wins a breeding site in a type-*t* patch; *ν*_t_ is the expected reproductive value of an individual in a type-*t* patch; *p*_m_ is the frequency of type-*m* patches in the population; *r*_it_ is either the relatedness of a rank-*i* mother in a type-*t* patch to one of her offspring (when dispersal is under maternal control), or else the relatedness of the offspring to itself (when dispersal is under offspring control); *h*_t_ is the probability that a random individual sampled after dispersal was born in the local patch (i.e. the probability of philopatry); *U*_jt_ is the probability that this philopatric individual was produced by the rank-*j* mother; and *ρ*_ijt_ is the relatedness of the rank-i mother (when dispersal is under maternal control) or an offspring of the rank-*i* mother (when dispersal is under offspring control) to an offspring of the rank-*j* mother in the same type-*t* patch (see Supporting Information for more details).

If dispersal is under maternal control, then *r*_it_ = *ρ*_iit_, as both of these quantities describe the relatedness of the rank-*i* mother to her own offspring. However, if dispersal is under offspring control, then *r*_i_ is the relatedness of the focal offspring to itself, whilst *ρ*_iit_ is its relatedness to its siblings. Condition (1) holds for both the asexual and sexual models, and also for haploid, diploid and haplodiploid modes of inheritance. Under the sexual reproduction model, the quantities described in condition (1) are sex-specific: for instance, if we are considering the dispersal of females, then *U*_jt_ is the probability that a random philopatric female is a daughter of a rank-j mother in a type-*t* patch.

### The constant philopater hypothesis

Of key interest is the quantity *N*_it_ = *N*_t_*U*_it_, which describes the number of philopatric offspring produced by a rank-i mother in a type-*t* patch, where *N*_t_ is the total number of philopatric offspring in the focal patch. Note that: the relatedness of a mother to her offspring, and the relatedness of the offspring to itself, are both independent of the mother’s rank, so we may write *r*_it_ = *r*_t_ for all i ∈ I, and all t ∈ T; the relatedness of an offspring to its siblings is independent of its mother’s rank, so we may write *ρ*_iit_ = *ρ*_t_ for all i ∈ I, and all t ∈ T; and the relatedness of a mother to another mother’s offspring, and the relatedness of an offspring to another mother’s offspring, is independent of the rank of either mother, so we may write *ρ*_ijt_ = P_t_ for all t ∈ T, all i ∈ I, and all j ∈ I, j ≠ i. Accordingly, ∑_*j∈I*_ *U*_*jt*_*ρ*_*ijt*_*= U*_*it*_*ρ*_*t*_ *+*P_*t*_ ∑_*j∈I,j ≠ i*_*U*_jt_*= U*_it_*ρ*_t_*+*P_t_(1–*U*_it_), and condition (1) can be rewritten as

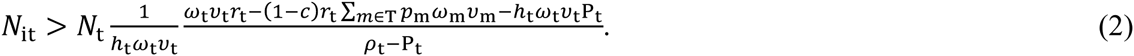

That is, the number of philopatric offspring produced by each rank-*i* mother in a type-*t* patch is favoured to converge upon the RHS of condition (2) and, because this quantity is independent of i, natural selection favours the number of philopatric offspring produced by each and every mother to converge upon the same number (i.e. *N*_it_ = *N*_t_*, and *U*_it_ = *U*_t_*), irrespective of the total number of offspring that she produces and her sex allocation. This result holds for both asexual and sexual reproduction under haploid, diploid, and haplodiploid inheritance, for female and/or male dispersal and for maternal or offspring control of dispersal. In analogy with the CMH, we term this invariant result the “constant philopater hypothesis” (CPH).

The CPH emerges from a balance between the mortality risk of dispersing and the kin-competition consequences of philopatry. From condition (1) we see that: because both the relatedness of a mother to her own offspring and also the relatedness of an offspring to itself are independent of maternal rank, the impact of the mortality cost of dispersal is the same for all mothers within each patch (-*r*_it_*ν*_t_+(1-*c*)*r*_it_∑_m∈T_*p*_m_*ν*_m_ = -*r*_t_*ν*_t_+(1-*c*)*r*_it_∑_m∈T_*p*_m_*ν*_m_ for all i ∈ I, and t ∈ T); because both the relatedness of a mother to another mother’s offspring and also the relatedness of an offspring to another mother’s offspring are independent of maternal rank (*ρ*_ijt_ = P_t_ for all t ∈ T, all i ∈ I, and all j ∈ I, j ≠ i), the offspring of all mothers experience the same strength of kin competition if all mothers produce the same number of philopatric offspring (*h*_t_*ν*_t_*U*_t_*\**(*r*_t_+(*n*-1)P_t_) under maternal control, or *h*_t_*ν*_t_*U*_t_*\**(*r*_t_+(*n*-1)*ρ*_t_) under offspring control); and, because any correlation that does arise between maternal rank and number of philopatric offspring leads to stronger kin competition among the offspring of mothers who produce more philopatric offspring, which favours such mothers to reduce their number of philopatric offspring, any correlation between rank and number of philopatric offspring will tend to disappear.

All mothers are favoured to produce the same number of philopatric offspring, but various constraints may interfere with their ability to do so. One possible constraint is that some low-ranking mothers are unable to produce the requisite number of philopatric offspring even if none of their offspring disperse, on account of their low fecundity. In this case, the CPH invariant breaks down, analogous to the breakdown of the CMH when some mothers are of such low fecundity that they cannot produce the requisite number of sons even if all of their offspring are male (Frank, 1985, 1987c).

### Within-patch heterogeneity

Above we have shown that the CPH holds under a very general set of assumptions, and we have expressed this result in terms of emergent quantities such as the relatedness and the probability of philopatry. Here we express these emergent quantities as a function of the underlying ecological and demographic parameters, which enables us to explicitly determine the optimal dispersal behaviour of offspring in particular scenarios. Here we focus on a particular case to illustrate how different model parameters mediate the optimal dispersal rates of offspring. We then contrast the optimal dispersal behaviour of offspring under maternal control with the optimal dispersal behaviour under offspring control to understand the role of the CPH in mediating parent-offspring conflict over dispersal.

We focus on a particular case in which there are two asexually-reproducing mothers per patch: a rank-1 mother with relatively-high fecundity (denoted by *F*_1_), and a rank-2 mother with relatively-low fecundity (denoted by *F*_2_). We denote the reproductive inequality between females by *s*, where *s* = 1-(*F*_2_/*F*_1_). We find that the probability of dispersal of offspring of high-fecundity mothers rises, whilst the probability of dispersal of offspring of low-fecundity mothers falls, as the reproductive inequality between mothers rises (Fig. 1). On the one hand, offspring of high-fecundity mothers and offspring of low-fecundity mothers both suffer the same cost of dispersal (*c*), and the relatedness between a focal offspring and herself is equal (*r*_1_ = *r*_2_ = 1), so the first term in inequality (1) is the same for both offspring (i.e. -*c r*_1_ = -*c r*_2_). But, on the other hand, all else being equal, the number of philopatric offspring of the high-fecundity mother is greater than that of the low-fecundity mother (*U*_1_ > *U*_2_): accordingly, the expected relatedness between a focal offspring of the high-fecundity mother and a random offspring in the patch is greater than the expected relatedness between a focal offspring of the low-fecundity mother and a random offspring in the patch (i.e. *h*(*U*_1_*ρ*_11_+*U*_2_*ρ*_12_) > *h*(*U*_1_*ρ*_21_+*U*_2_*ρ*_22_), where *ρ*_11_ = *ρ*_22_ = 1, and *ρ*_12_ = *ρ*_21_ = *ρ*). Therefore, the selection pressure for dispersal of offspring of high-fecundity mothers is stronger than the selection pressure for dispersal of offspring of low-fecundity mothers.

**Figure 1.**
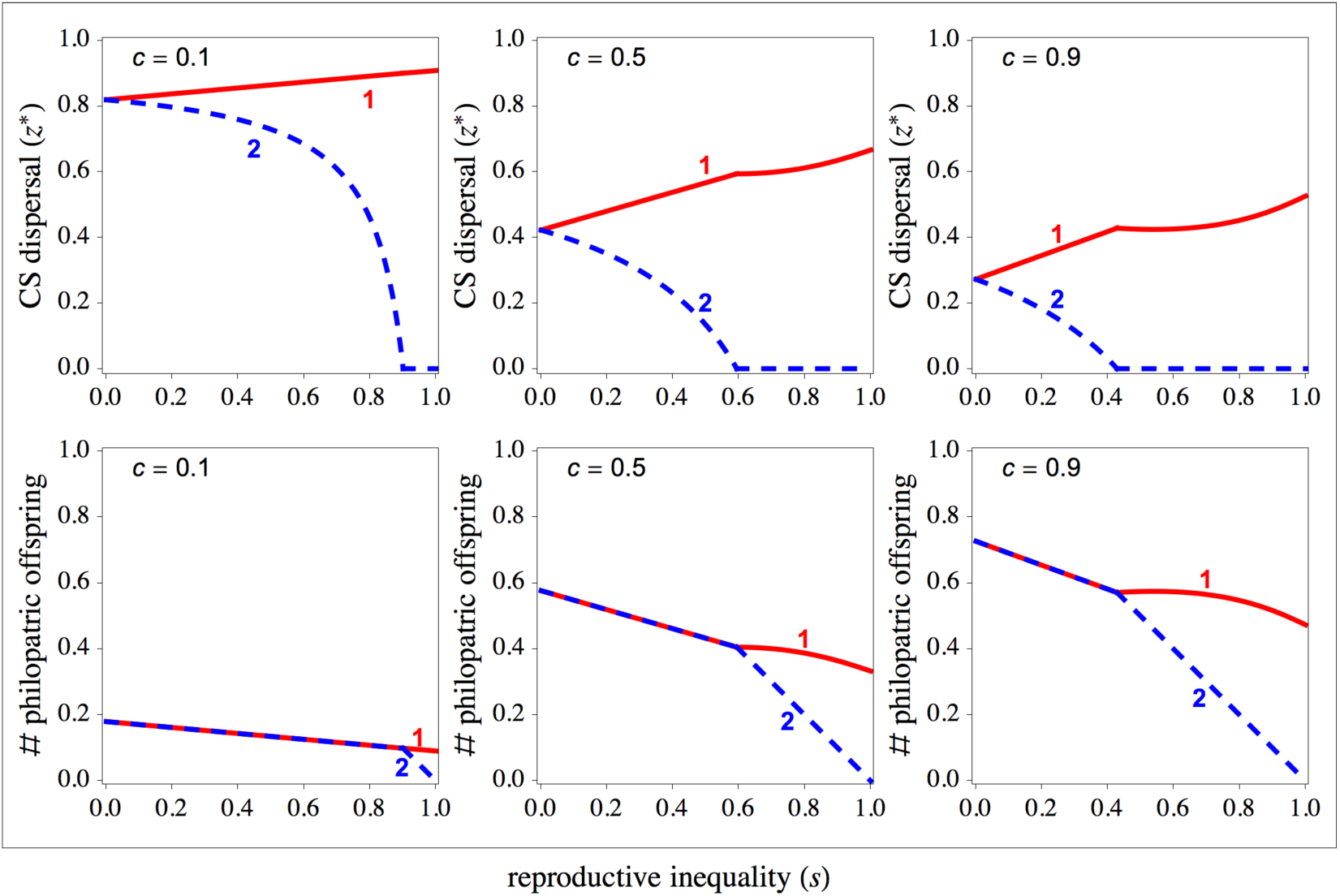
Convergence stable dispersal rates in heterogeneous groups. The CS dispersal strategies of offspring of high-fecundity rank-1 breeders (*z*_1_*, solid lines) and of offspring of low-fecundity rank-2 breeders (*z*_2_*, dashed lines) as a function of the reproductive inequality (*s*) for varying cost of dispersal (*c*). The dispersal rate of offspring of high-fecundity breeders is greater than that of offspring of low-fecundity breeders (i.e. *z*_1_* > *z*_2_*). All breeders produce the same number of offspring that remain in the natal patch as long as low-fecundity mothers give birth to a sufficiently high number of offspring.

We also find that the mean probability of dispersal falls as the cost of dispersal rises (Fig. 1). As the cost of dispersal rises, the first term in inequality (1) decreases and the second term in inequality (1) increases. As the effect on the first term is stronger than the effect on the second term, the overall effect of increasing the cost of dispersal is that dispersal becomes less evolutionarily advantageous.

The number of philopatric offspring of the high-fecundity mother rises as the reproductive inequality between the two mothers increases, and as the cost of dispersal increases. So long as this number is not too high, low-fecundity mothers are able to match it (i.e. 1-*z*_1_* = (1-*s*)(1-*z*_2_*)). However, if the number of philopatric offspring of high-fecundity mothers is too high (due to high *s* and / or high *c*), then low-fecundity mothers cannot produce the requisite number of philopatric offspring even if none of their offspring dispersal, and in such scenarios the CPH breaks down (Fig. 1).

### Between-patch heterogeneity

#### Temporally-stable environments

We now consider a heterogeneous population in which there are type-1 patches with high resource-availability and type-2 patches with low resource-availability. We define the reproductive inequality between patches as *s*_b_ = 1-(*F*_12_/*F*_11_), and the reproductive inequality within patches as *s*_1_ = 1-(*F*_21_/*F*_11_) = *s*_2_ = 1-(*F*_22_/*F*_12_) = *s*. We first consider a spatially-heterogeneous environment in which patches retain their type over generations. We find that the average probability of dispersal is higher from low-quality type-2 patches than from high-quality type-1 patches (Fig. 2, panel (c)). As a result, high-quality patches have more non-dispersing offspring than low-quality patches. However, in both types of patches, higher-ranking mothers disperse more offspring than lower-ranking mothers, and, as long as inequality within patches is sufficiently small, both high- and low-rank mothers produce exactly the same number of philopatric offspring irrespective of the quality of their patch (Fig. 2, panel (f)).

**Figure 2.**
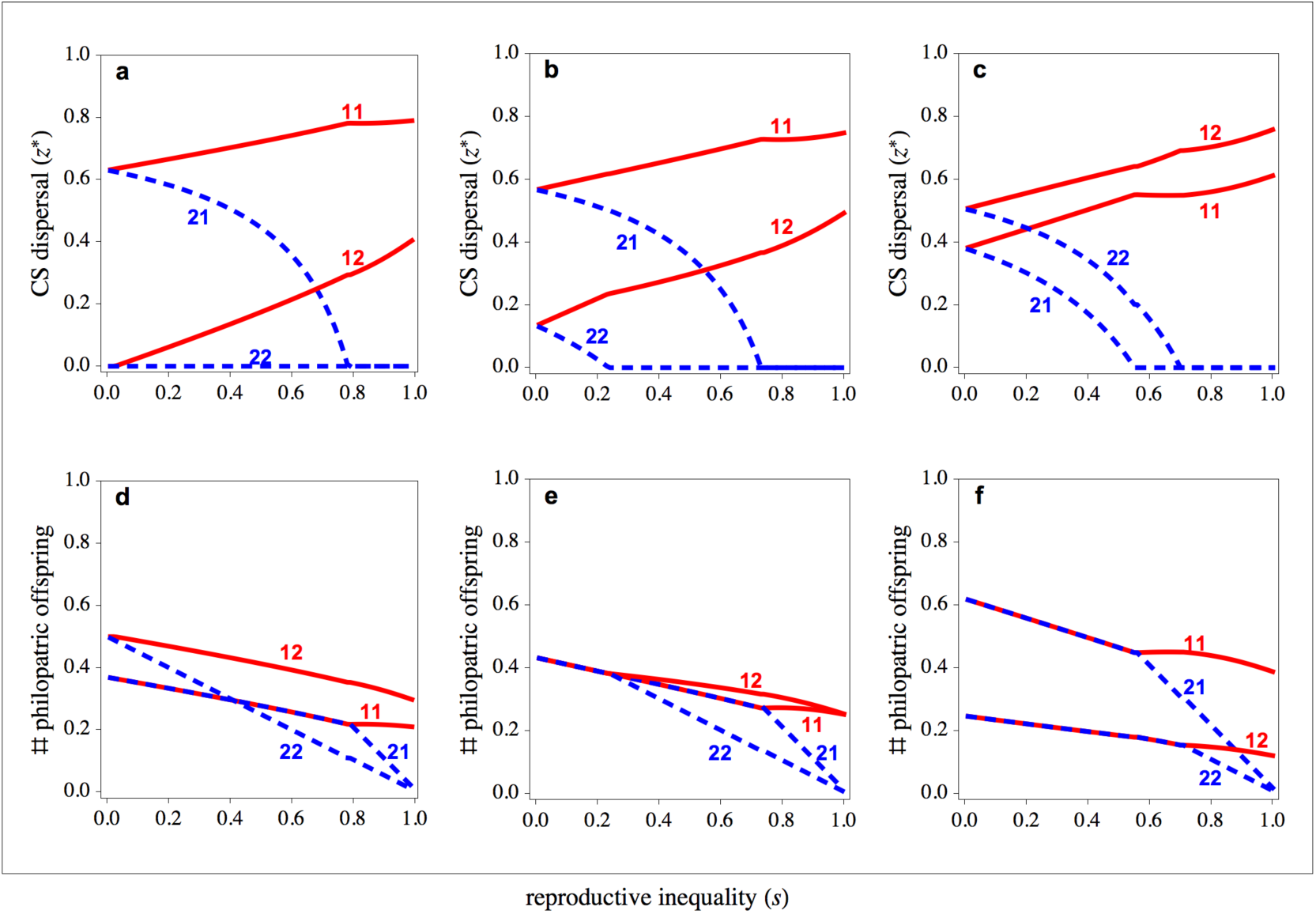
Convergence stable dispersal rates in heterogeneous populations. The CS dispersal strategies of offspring of high-fecundity rank-1 breeders (*z*_1X_*, solid lines) and of offspring of low-fecundity rank-2 breeders (*z*_2X_*, dashed lines) in high resource-availability rank-1 patches (*z* *) and in low resource-availability rank-2 patches (*z* *) as a function of the reproductive inequality (*s*) for temporally stable, unpredictable, and seasonal environments. (a,d) In temporally seasonal environments average dispersal is higher from rank-1 patches, and the CPH holds as long as inequality is sufficiently small. (b,e). In temporally unpredictable environments average dispersal is higher from rank-1 patches, and the CPH holds both within and between patches as long as inequality is sufficiently small. (c,f) In temporally stable environments average dispersal is higher from rank-2 patches, and the CPH holds as long as inequality is sufficiently small. Parameter values: *c* = 0.50, *p* = 0.50.

#### Temporally-unpredictable environments

We next consider unpredictable environments in which a patch’s type in the next generation is independent of its type in the current generation. Under such circumstances, the expected reproductive value is identical across patches. Thus, *ν*_t_ = *ν*, for all t ∈ T. Moreover, the relatedness coefficients are also identical across patches. Thus, *r* = *r*_t_, *ρ*_t_ = *ρ*, and P_t_ = P. Therefore, inequality (2) becomes

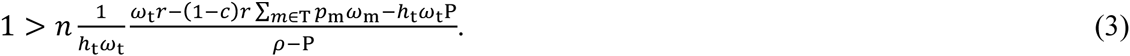

This means that, at equilibrium, *h*_t_* = *h*\* and *ω*_t_* = *ω*\*, and therefore natural selection favours the number of philopatric offspring produced by each and every mother to converge upon the same number (i.e. *N*_it_ = *N*_t_* = *N*\*, and *U*_it_ = *U*_t_* = *U*\*). Thus, in unpredictable environments, the CPH holds not only within each patch, but also between patches (Fig. 2, panel (e)).

#### Seasonal environments

Finally, we consider seasonal environments in which a patch always changes its type from one generation to the next. We find that the average probability of dispersal is higher from high-quality type-1 patches than from low-quality type-2 patches (Fig. 2, panel (a)). As a result, low-quality patches have more philopatric offspring than high-quality patches. However, in both types of patches, higher rank mothers disperse more offspring than lower rank mothers, and, as long as inequality within patches is sufficiently small, both high- and low-rank mothers produce exactly the same number of philopatric offspring irrespective of the quality of their patch (see Fig. 2, panel (d)).

### Parent-offspring conflict

Although the CPH result obtains irrespective of whether dispersal is controlled by the offspring themselves or by their mother, we find that the level of dispersal that is favoured does depend upon whose control it is under. This recovers Motro’s (1983) result that an evolutionary conflict of interest often exists between mother and offspring with regards to dispersal, with mothers generally preferring that their offspring disperse at a rate that is higher than the rate at which the offspring would prefer to disperse themselves. This is on account of the mother being equally related to those offspring that disperse and their siblings that benefit from the resulting relaxation of kin competition, and her offspring being more related to themselves than they are to each other (see also Frank, 1986; Taylor, 1988b; Gandon, 1999; Starrfelt & Kokko, 2010).

Our model has crucially incorporated heterogeneity in maternal condition, and this allows us to investigate how such heterogeneity mediates the parent-offspring conflict of interests with respect to dispersal. Here, we determine whether the potential for conflict is greater in families with more resources (i.e. families with high-fecundity rank-1 mothers) or fewer resources (i.e. families with low-fecundity rank-2 mothers). We consider two scenarios: one in which offspring have complete information about their mothers’ rank (i.e. conditional dispersal); and one in which offspring have no information about their mothers’ rank (i.e. unconditional dispersal). We first focus on cases in which offspring have complete information about their mothers’ rank. Here, we find that mothers always prefer greater dispersal rates of offspring than the offspring, irrespective of the resources available for each family (Fig. 3). However, the difference between the optimal behaviour from the mother’s perspective and the optimal behaviour from the offspring’s perspective is not the same for the different types of families. In particular, we find that for lower inequality, conflict is more pronounced within resource-poor families than within resource-rich families (Fig. 3). As inequality between families rises the optimal dispersal rate of offspring in resource-rich families rises, whereas the optimal dispersal rate of offspring in resource-poor families falls, irrespective of who controls the dispersal rate of offspring. When the inequality between families is sufficiently large, resource-poor families hit a threshold beyond which all their offspring are philopatric, independently of who controls the dispersal rate of offspring. At this point the conflict within resource-poor families ceases, whilst it still exists within resource-rich families (Fig. 3). In summary, when inequality is low, resource-poor mothers suffer more parent-offspring conflict over offspring dispersal than resource-rich families, but they still produce a fair amount offspring. When inequality is high, there is less conflict within resource-poor families, but their fecundity is very low.

**Figure 3:**
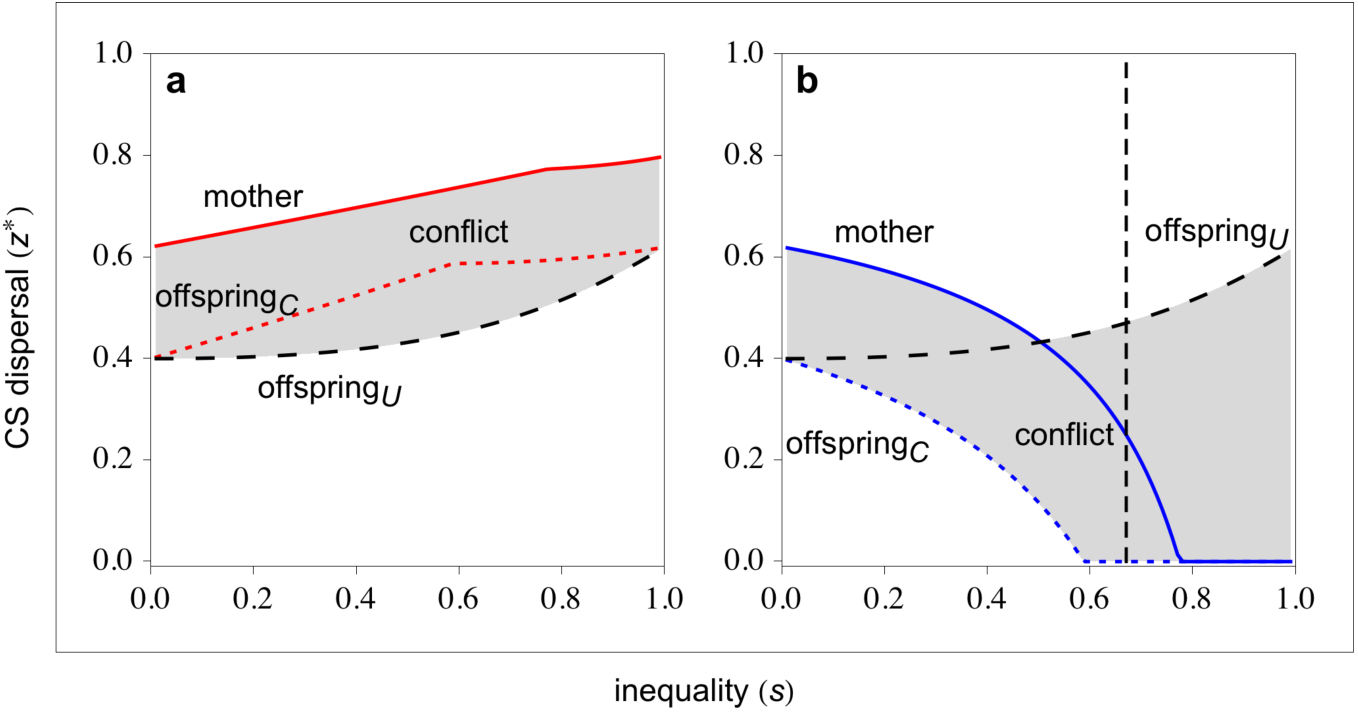
Parent-offspring conflict. The CS dispersal strategies of mothers (solid lines) and of daughters under complete maternity information (offspring_C_, dotted lines) and under no maternity information (offspring_U_, dashed lines) for (a) rank-1 resource-rich families, and (b) rank-2 resource-poor families. Under lower reproductive inequality, parent-offspring conflict is more intense for low-fecundity families. For resource-rich families, parent-offspring conflict is more intense under no-maternity information irrespective of the inequality between families. For resource-poor families, parent-offspring conflict is less intense under no-maternity information to the left of the vertical dashed line. The number of philopatric offspring is in arbitrary units. Parameter values: *c* = 0.25.

We next contrast cases in which offspring have complete information about their mothers’ rank with cases in which offspring have no information about their mothers’ rank. This allows us to investigate the circumstances under which mothers are selectively favoured to inform their offspring as to their rank versus withholding this contextual information. We find that when offspring know that they have rank-1 mothers, parent-offspring conflict is less strong than when offspring do not know the rank of their mothers (Fig. 3, panel (a)). This suggests that rank-1 mothers should disclose full information about their status to their offspring in order to minimise parent-offspring conflict. In contrast, we find that when offspring know that they have rank-2 mothers, parent-offspring conflict is stronger than when offspring do not know the rank of their mothers, as long as inequality is sufficiently small (Fig. 3, panel (b)). This suggests that rank-2 mothers should withhold information about their status from their offspring in order to minimise parent-offspring conflict. These conflicting selective forces generate an informational battleground between rank-1 and rank-2 mothers, in which rank-1 mothers are favoured to disclose maternity information to offspring in the group whilst rank-2 mothers are favoured to withhold it.

### Allomaternal control of dispersal

Above, we have considered that control of offspring dispersal occurs either by the offspring themselves or by their mothers. Whilst this may often be the case, in other situations mothers may control the dispersal traits of offspring other than their own. This may be particularly important when differences in fecundity between mothers are also extended to other behavioural traits such as dominance over other group members. First we consider a case in which the high-fecundity breeder has full control over the dispersal of her own offspring, but varies in the degree of control, denoted by *α*, over the offspring of the low-fecundity mother, with 0 ≤ *α* ≤ 1. We find that the CPH holds as long as the high-fecundity mother does not exert any control over the dispersal of the low-fecundity mother’s offspring (i.e. when *α* = 0; Fig. 4). However, when the degree of control by the high-fecundity mother increases, the dispersal probability of their own offspring decreases, whilst the dispersal probability of the low-fecundity mother’s offspring increases (Fig. 4, panel(a)). Indeed, when the high-fecundity mother reaches a certain degree of control, all of the low-fecundity mother’s offspring are forced to disperse (i.e. *z*_2_ = 1). We obtain similar results when we allow the low-fecundity mother control the dispersal of the high-fecundity mother’s offspring, where we denote the degree of control of the low-fecundity mother by *β*. When the degree of control by the low-fecundity mother increases, the dispersal probability of their own offspring decreases, whilst the dispersal probability of the low-fecundity mother’s offspring increases (Fig. 4, panel(b)). If the degree of control is sufficiently high, all offspring of high-fecundity rank-1 mothers are forced to disperse, whilst all offspring of low-fecundity rank-2 mothers remain in the local patch. When the low-fecundity mother has no control over the high-fecundity mother’s offspring (i.e. when *β* = 0), the CPH holds, but not otherwise (i.e. when *β* > 0; Fig. 4).

**Figure 4.**
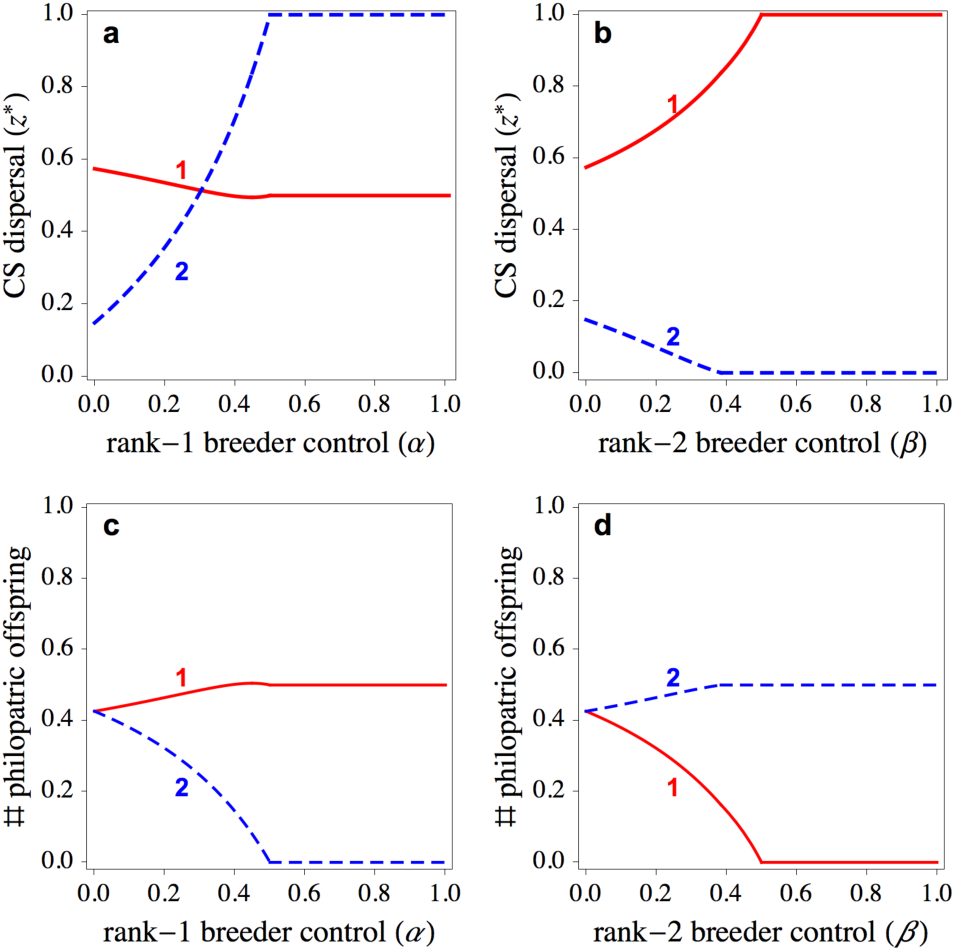
Allomaternal control of dispersal. The CS dispersal strategies of offspring and the number of philopatric offspring as a function of rank-1 high-fecundity and rank-2 low-fecundity mothers degree of control. When mothers control the dispersal of their own offspring (i.e. *α* = 0, and *β* = 0) the CPH holds. (a,c) When high-fecundity mothers increase their control over the dispersal of low-fecundity mothers’ offspring, the dispersal of low-fecundity mothers’ offspring rises whilst the dispersal of their own offspring falls. (b,d) When low-fecundity mothers increase their control over the dispersal of high-fecundity mothers’ offspring, the dispersal of high-fecundity mothers’ offspring rises whilst the dispersal of their own offspring falls. (c,d) The CPH breaks down when mothers do not control the dispersal of their own offspring or when offspring do not control their own dispersal. The number of philopatric offspring is in arbitrary units. Parameter values: *c* = 0.25, *s* = 0.5.

### The CPH underpins a family of invariance results

To the extent that any trait may be coincident with an individual’s dispersal status, the CPH underpins a whole family of invariance results. For example, if dispersing individuals engage in aggressive behaviour whilst non-dispersing individuals are more docile (e.g. El Mouden & Gardner, 2008), then the present CPH result could be reframed as a “constant non-aggressor hypothesis”. The important caveat here is that such derivative invariants are only expected to hold insofar as the focal trait is tightly coupled to dispersal status, and the fact that incomplete coupling leads to a failure of these invariants whilst the CPH continues to hold confirms that the CPH is the more fundamental invariant.

One such derivative invariant that has been previously described is the “constant female hypothesis” (CFH; Frank, 1987c, 1998). This is concerned with “local resource competition” (Clark, 1978) scenarios in which females are philopatric and males are the dispersing sex, and the CFH predicts that more fecund mothers will invest relatively less into daughters than will less fecund mothers, such that all mothers will produce the same number of daughters, irrespective of their fecundity. This is because the selection gradient acting on the sex allocation strategy shows properties that are identical to those of the selection gradient acting on dispersal. Namely, if we assume that the sex-ratio of a mother (i.e. *σ*_it_) is now an evolving trait, rather than a parameter, the selection gradient acting on the sex allocation strategy of a mother is given by

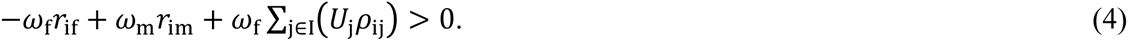

As in the CPH, the relatedness coefficients are independent of the mother’s rank. Thus, *r*_i_ = *r* and *ρ*_ii_ = *ρ* for all i ∈ I; *ρ*_ij_ = P for all i ∈ I, and all j ∈ I, j ≠ i. Thus, mothers adjust their sex ratio such that each and every mother converge upon the same number of daughters (i.e. *N*_i_ = *N*\*, and *U*_i_ = *U*\*).

However, this invariant result only holds when all daughters are philopatric. If females exhibit at least some propensity to disperse, then the selection gradient acting on the sex ratio is given by

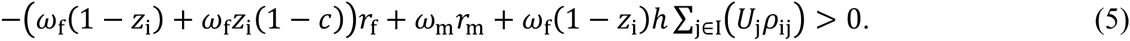

This means that the first term of the selection gradient now depends on the fecundity of the focal mother, and therefore the CFH no longer holds. The CPH, by contrast, does hold, irrespective of the sex ratio produced by each mother. That is, it is the CPH that underpins the CFH, and not the reverse.

## Discussion

We have described a new life-history invariant result for dispersal evolution. Specifically, we have found that natural selection favours all mothers to produce the same number of philopatric offspring, irrespective of variation between mothers in the total number of offspring that they produce. This is because kin competition, arising from a failure to disperse, is related to the number, rather than the proportion, of a mother’s philopatric offspring. In analogy with the similar “constant male hypothesis” (CMH) of the sex allocation literature (Frank, 1987c, 1998), we term this result the “constant philopater hypothesis” (CPH).

Such invariance results provide testable predictions in their own right, and also promote the interplay of theory and empirical testing by reducing the extent to which extraneous genetic, ecological and demographic parameters are confounding in comparative analyses (e.g. West et al., 2001; Rodrigues & Gardner, 2015). Moreover, they also facilitate the development and conceptualisation of theory. For example, the invariant relationship between helping and harming, on the one hand, and degree of population viscosity, on the other hand (Taylor, 1992; El Mouden & Gardner, 2008), has been used to demonstrate that heterogeneity in resource availability per se – and not any conflating effect of viscosity itself – modulates the evolution of helping and harming in viscous populations (Rodrigues & Gardner, 2012, 2013a). The CPH invariance prediction is readily amenable to empirical testing, as it is robust to variation in difficult-to-measure quantities such as the mortality risk associated with dispersal. Social groups in different species often comprise multiple breeders that vary in their fecundity (reproductive skew), and in some cases there is variation in the proportion / number of dispersers produced by each breeder (e.g. Crespi & Taylor 1990; Innocent et al., 2010). Our theory predicts that dispersal rates (or the fraction of dispersal morphs) should be higher for more productive breeders, and that at the same time the number of philopatric offspring should be equal for each breeder.

In terms of reaction-norms, the CPH means that mothers with fecundity below a certain threshold should produce no dispersing offspring, while mothers with fecundity above that threshold should exhibit a positive correlation between their fecundity and the dispersal rate of their offspring. Such reaction norms with a critical threshold have also been observed in the context of the evolution of dispersal conditional on the overall number of individuals in a patch (Crespi & Taylor, 1990; Ezoe & Iwasa, 1997; Kisdi, 2004; Rodrigues & Johnstone, 2014). Under certain conditions, this means that differences in density between patches before dispersal are eroded after dispersal. Specifically, we have shown that the CPH holds both within and between patches when the environment is temporally unpredictable but not for other types of temporal variation. This implies that there are two forces mediating the evolution of dispersal: one acting between patches that tends to equalise or enhance differences in density between them; and one acting within patches that tends to equalise differences in number of philopatric offspring among group members. These two forces may be operating simultaneously in natural population, and future empirical studies should take both into consideration.

We have shown that the adaptive adjustment of offspring dispersal conditional on maternal fecundity may have a dramatic impact on the amount of kin competition that each offspring experiences. More specifically, this means that variation in fecundity among breeders is not translated into an equivalent variation in kin competition among offspring. Indeed, owing to the CPH, the amount of kin competition may be precisely the same, irrespective of a mother’s fecundity. This has wide-reaching implications for the evolution of social behaviour within groups. We have shown how the CPH underlies the “constant female hypothesis”, an invariant result that has been previously described in the sex allocation literature (Frank, 1987c, 1998). Another topic for which the CPH may have important implications is reproductive skew, which has been shown to promote the evolution of harming by high-fecundity mothers and helping by low-fecundity mothers (Johnstone, 2008). Crucially, that result has been derived under the assumption that, while helping and harming are conditional on a mother’s fecundity, dispersal of offspring is not. An immediate consequence of the CPH is that, if offspring disperse conditionally, according to maternal fecundity, the asymmetry in the level of kin competition between high- and low-fecundity mothers vanishes, such that helping and harming are no longer favoured. This suggests a promising avenue for future theoretical and empirical study.

We have also shown that there will typically be a conflict of interests between parent and offspring with respect to the latter’s probability of dispersing, and that the intensity of such conflict is modulated by heterogeneity in parental condition, and hence is liable to vary between families. We find that if inequality in fecundity is sufficiently low, the intensity of parent-offspring conflict is greater in resource-poor families but, by contrast, if the inequality is sufficiently high, the conflict within resource-poor families may vanish, with parents and offspring agreed that there should be no dispersal. To the extent that within-family conflict has a negative impact on a mother’s fecundity, this result suggests that parent-offspring conflict may either reinforce inequality between families (when inequality is relatively low) or may attenuate inequality between families (when inequality is relatively high).

On account of our finding that parent and offspring dispersal optima depend upon the degree of heterogeneity in fecundity across families, we have uncovered a new informational battleground over dispersal, with high-fecundity mothers being favoured to disclose full information about their status to all the offspring in the group, and low-fecundity mothers being favoured to withhold this information. The resolution of this informational conflict will depend upon the specific biology of particular species (for reviews see Godfray, 1995; Kilner & Hinde, 2008). There are many examples of mothers disclosing contextual information to their offspring: in daphnia, for instance, mothers provide accurate information about the presence of predators in the local environment, and offspring respond to this information by developing a protective helmet (Tollrian & Dodson, 1999). Conversely, there are examples of mothers withholding information or actively deceiving their offspring with regards to the circumstances in which they find themselves: in black-headed gulls, *Larus ridibundus*, for instance, mothers appear to adjust yolk androgen concentration in eggs in order to manipulate the offspring’s perception of their birth order in the brood (Eising et al., 2001).

More generally, we suggest that the resolution of this informational conflict will depend on whether mothers are: (i) constrained to either honestly communicate their rank to their offspring or else withhold this information; or (ii) able to honestly communicate, withhold the information or deceive their offspring with regards to their rank. If deception is not an option, then in this simple binary scenario an offspring will always be able to correctly determine her mother’s rank, either because her mother honestly communicates the fact that she is of rank-1 or else because her mother communicates no information, which enables the offspring to infer that she is of rank-2, and this system of signalling will be stably maintained by the coincidence of interests of the rank-1 mother and her offspring. However, if unconstrained deception is an option, then all mothers are expected to communicate that they are of rank-1, which provides no useful information to their offspring, and hence this system of communication is expected to collapse. The resolution of this conflict represents a further avenue for future research.

Our model provides an explanation for different patterns of dispersal within social groups depending on the degree of control by each group member, which can change the sign of rank-dependent dispersal. If each mother controls the dispersal of their own offspring or if offspring control their own dispersal, then we should expect a positive correlation between mother’s fecundity and offspring propensity to disperse – i.e. positive rank-dependent dispersal. Under allomaternal control of offspring dispersal the mother with a higher degree of control is expected to force offspring of other mothers to disperse and therefore their own offspring are less likely to disperse. If, for instance, the dominant mother controls the dispersal of offspring in the social group, then we should expected negative rank-dependent dispersal. For example, in meerkats the dominant is more likely to force distantly related offspring to disperse than their close relatives, and therefore offspring of lower rank mothers are more likely to disperse than offspring of higher rank mothers (i.e. negative rank-dependent dispersal rates; Clutton-Brock et al., 2010). By contrast, in the red-fronted lemurs, there is no correlation between dispersal and kinship, and therefore we should not expect negative rank-dependent dispersal rates (Kappeler & Fichtel, 2012, reviewed in Clutton-Brock, 2013). More generally, while in our model we have considered a simple control parameter, more species-specific resolution models can be adopted (e.g. Godfray, 1995; Kilner & Hinde, 2008). These possibilities also represent avenues for future theoretical and empirical exploration.

The CPH result emerges from key symmetries in relatedness, for instance the independence of the relatedness between two mothers breeding in the same patch with respect to their rank and hence their share of the group’s total fecundity. This situation obtains in the present model owing to our assumption that rank is not inherited. However, more generally, rank may be heritable, to some extent, such that high-ranking females tend to be the daughters of highly-fecund mothers, in which case they may be more likely to breed alongside sisters than are females of lower rank, which could lead to a positive correlation between rank / fecundity and relatedness to group mates. Alternatively, whilst we have considered the cost of dispersal to be paid in terms of mortality, dispersal may also incur fecundity costs, which again could lead to a positive correlation between fecundity and relatedness, owing to low-fecundity dispersers being unrelated to their group mates.

Finally, although our results hold under a wide range of model assumptions, we have not studied the effects of many other potentially-relevant factors. It is likely that, in some of these cases, our model will fail to conform to empirical data. However, by highlighting those scenarios in which our model’s key assumptions are not met, our result may be used to illuminate otherwise obscured biological details, concerning a species’ ecology, demography, phenotypic plasticity or cognition. In this respect, the model also establishes a baseline scenario, which may help to understand and interpret new empirical data and future mathematical results.

## Acknowledgements

We thank: Rebecca Kilner for encouragement and support; and the Behavioural Ecology Group, Cambridge, for helpful discussion. This research was supported by Wolfson College Cambridge (A.M.M.R.) and the Natural Environment Research Council (grant no. NE/K009524/1) (A.G.).

